# Cdc42 negatively regulates endocytosis during apical plasma membrane maintenance and development in mouse tubular organs *in vivo*

**DOI:** 10.1101/402677

**Authors:** Akiko Shitara, Lenka Malec, Seham Ebrahim, Desu Chen, Christopher Bleck, Matthew P. Hoffman, Roberto Weigert

## Abstract

Lumen establishment and maintenance are fundamental for tubular organs physiological functions. Most of the studies investigating the mechanisms regulating this process have been carried out in cell cultures or in smaller organisms, whereas little has been done in mammalian model systems *in vivo*. Here we used the salivary glands of live mice to examine the role of the small GTPase Cdc42 in the regulation of the homeostasis of the intercellular canaliculi, a specialized apical domain of the acinar cells, where protein and fluid secretion occur. Depletion of Cdc42 in adult mice induced a significant expansion of the apical canaliculi, whereas depletion at late embryonic stages resulted in a complete inhibition of their post-natal formation. In addition, intravital subcellular microscopy revealed that reduced levels of Cdc42 affected membrane trafficking from and towards the plasma membrane, highlighting a novel role for Cdc42 in membrane remodeling through the negative regulation of selected endocytic pathways.

## Introduction

Epithelial cells have two specialized surface domains, the basolateral and the apical, each with a distinct composition and function (Bryant and Mostov, 2008; Willenborg and Prekeris, 2011). In tubular organs such as the salivary and mammary glands, lung, kidney, pancreas and intestine, the apical plasma membrane (APM) forms the lumens. These specialized areas of the cell are implicated in extensive protein, fluid, and electrolyte secretion and uptake, and therefore undergo constant remodeling (Masedunskas et al., 2011b; Tepass, 2012; Willenborg and Prekeris, 2011). The establishment and maintenance of the APM are regulated by a complex signaling cascade controlled by the small GTPase Cdc42 (Etienne-Manneville, 2004; Melendez et al., 2011). In its GTP-bound form, Cdc42 binds and activates PAR6, a member of the apical PAR polarity complex (PAR3–PAR6–aPKC) which together with the Crumbs (Crb– PALS–PATJ) and Scribble (Scrib–Dlg–Lgl) complexes dictate the positioning of the apical-basolateral border through the assembly of tight and adherens junctions (Bryant and Mostov, 2008). In addition, the polarity complexes regulate a series of downstream effectors, which coordinate the activation of both actin cytoskeleton and membrane trafficking, and are required for establishing and maintaining cell polarity (Etienne-Manneville, 2004; Melendez et al., 2011).

The role of Cdc42 in regulating the homeostasis and establishment of the APM has been extensively investigated in Madin-Darby canine kidney (MDCK) cells grown in purified extracellular matrix components (Bryant et al., 2010; Jaffe et al., 2008; Martin-Belmonte et al., 2007). The versatility of this experimental model has allowed the identification of a large number of molecules implicated in this process, and to tease out several details of this complex molecular machinery (Apodaca, 2010; Bryant and Mostov, 2008; Etienne-Manneville, 2004). In addition, substantial work has been performed in smaller multicellular organisms such as *Drosophila, C. Elegans*, and zebrafish (Balklava et al., 2007; Georgiou et al., 2008; Kamei et al., 2006; Pirraglia et al., 2010). However, only a limited amount of work has been carried out in mammalian organisms (i.e. rodents). These studies, which took advantage of the ablation of Cdc42 in specific adult tissues such as liver (van Hengel et al., 2008), pancreas (Kesavan et al., 2009), intestine (Melendez et al., 2013) and inner ear (Ueyama et al., 2014), showed a fundamental role of Cdc42 in regulating apical polarity, although they did not focus on the characterization of its role in regulating membrane remodeling and trafficking.

Here, we used a combination of intravital subcellular microscopy (ISMic), a light microscopy-based technique that enables imaging the dynamics of intracellular structures in live animals (Masedunskas et al., 2011a; Masedunskas et al., 2013; Milberg et al., 2014; Pittet and Weissleder, 2011; Weigert et al., 2013), and indirect immunofluorescence, to examine the role of Cdc42 in controlling maintenance and formation of the intercellular canaliculi (IC) in the acinar cells of the submandibular salivary glands (SMGs) in live mice. These structures are narrow tubes, formed by the APM of two adjacent acinar cells (Fig.1), which constitute a network spanning throughout the secretory acini (Fig. 1A) and connected to a central canaliculus that leads to the intercalated ducts (Tamarin and Sreebny, 1965). The IC play a fundamental role in the physiology of the salivary glands since they are the site where protein and fluid secretion occur. Cdc42 was ablated in either adult mice or on embryonic day 15. We found that reduction in the levels of Cdc42 in adult mice resulted in the involution of the IC, which progressively retracted from the basal border and underwent significant expansions. On the other hand, in neonatal mice, the lack of Cdc42 resulted in the complete inhibition of the formation of the IC. In parallel, we observed a massive stimulation of endocytic trafficking, thus suggesting a scenario in which Cdc42 negatively regulates endocytosis during maintenance and establishment of membrane polarity.

**Figure 1.**
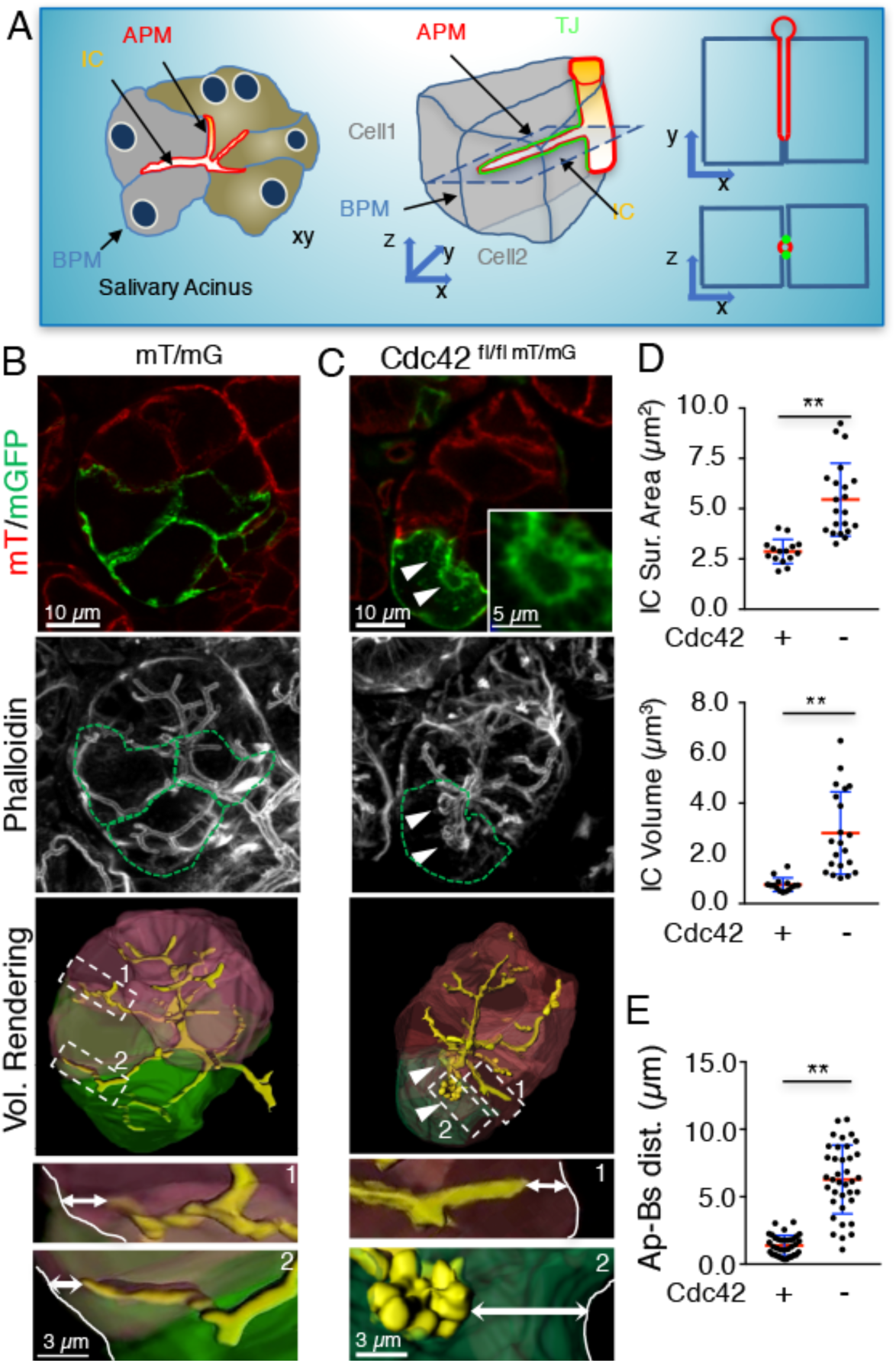
Cdc42-depletion induces the expansion of the intercellular canaliculi in adult mice. **A.** Diagram of the salivary glands acinus showing the structure of the IC formed by the APM of two adjacent cells. XY and XZ view of the canaliculi. Highlighted in green are the TJ. **B-C**. mT/mG (B) or Cdc42 ^fl/fl-mT/mG^(C) mice were transfected with adeno-Cre. After 3 weeks the glands were excised, labelled with Alexa 647-Phalloidin and imaged by confocal microscopy, as described in Material and Methods. Dashed green lines highlight the Cre-expressing cells. Arrowheads highlight the expanded ICs. Lower panels feature the volume rendering of representative acini (Cre-positive cells highlighted in green, Cre-negative cells highlighted in red) (See also Movie S1 and S2. Double arrows highlight the distance between the tip of the IC and the basal PM. **D**. Quantification of surface area and volume of IC in Cdc42 ^fl/fl-mT/mG^ mice. Cre-negative (Cdc42+) cells: Surface Area: 2.87 ± 0.59 μm^2^; Volume: 0.75 ± 0.27 μm^3^, N=15) and Cre-positive cells (Cdc42-) (Surface Area: 5.45 ± 1.81 μm^2^, Volume: 2.81 ± 1.64 μm^3^, N=21). Data are Means ± S.D.; N=number of IC from 11 acinar cells scored in 4 animals; ** p<0.01, unpaired Student’s t-test. **E**: Quantification of distance between the tip of the IC and the basal PM in Cre-negative (Cdc42+) (1.38 ± 0.75 μm, N=36) and Cre-positive (Cdc42-) (6.28 ± 2.55 μm, N=37). Data are Means ± S.D.; N=number of IC from 11 acinar cells from 5 animals; ** p<0.01, unpaired Student’s t-test.

## Results & Discussion

### Cdc42-depletion causes the expansion of the intercellular canaliculi in adult mice

In order to define the role of Cdc42 in maintaining the homeostasis of luminal structures in live adult mice, we used a Cre/loxP approach to deplete Cdc42 from the SMG acinar cells. Cdc42 floxed mice (Chen et al., 2006) were crossed with the Cre-recombinase (Cre) reporter mouse strain Rosa^mT/mG^ (mTmG) (Muzumdar et al., 2007), as shown in Fig. S1A (Cdc42 ^fl/fl-mT/mG^). The mT/mGFP probe provides a marker for both Cre expression and visualization of the plasma membrane (PM), as previously described (Masedunskas et al., 2011a; Milberg et al., 2017). Cre expression was induced in 5-10% of the acinar cells by the injection of a Cre-expressing adenovirus (Adeno-Cre) into the submandibular main duct (Milberg et al., 2017; Sramkova et al., 2009). In Cdc42 ^fl/fl-mT/mG^ mice, 3-4 weeks after the adenovirus injection, Cre-expressing cells exhibited large vacuolar structures labeled with membrane-targeted GFP (mGFP) and contiguous to the APM (Fig. 1C upper panel, arrowheads, and inset). F-actin labeling, which highlights the APM, revealed that these structures were the result of IC expansion (Fig. 1C middle panel, arrowheads, and inset). We also found that the expanded IC were primarily observed in cells with lower levels of Cdc42, as assessed by immunofluorescence (Fig. S1B). Z-stacks and volume rendering of the acini further confirmed that the APM bulged towards the interior of Cdc42-depleted cells (Fig. 1C, lower panel, and Movie S1). In contrast, no changes were observed in either Cre-negative cells (Fig. 1C), or Cre-positive cells in mTmG mice, that were used as additional controls (Fig. 1B, and Movie S1). Quantitative analysis showed that Cdc42-depletion resulted in the: 1) increase of both surface area and volume of the APM with respect to control cells (Fig. 1D); and 2) shortening of the IC length (Fig. 1E), as shown by measuring the distance between the tip of the IC and the basal PM, which suggests a repositioning of the apical-lateral border (Fig. 1B and 1C lower panels). Interestingly, the apical-lateral border was altered regardless it was shared between one or two cells lacking Cdc42 (Fig. S1C). Finally, depletion of the small GTPases RhoA and Rac1 did not have any effect on the morphology of the IC, thus suggesting a specific role for Cdc42 in their maintenance (Fig. S1D).

### Cdc42-depletion leads to loss of PAR6 and F-actin at the APM

We reasoned that changes in the morphology of the IC in Cdc42-depleted cells could be the result of a defect in the maintenance of cell polarity. Therefore, we investigated the recruitment of the Par polarity complex on the salivary glands APM. In cell culture and in small organisms this complex has been shown to regulate various functions such as, establishment and maintenance of polarity, cytoskeletal assembly, tight junction (TJ) formation, and positioning of the apical–lateral border (Morais-de-Sá et al., 2010; Suzuki and Ohno, 2006; Tepass, 2012; Walther and Pichaud, 2010). We found that in the IC of Cdc42-depleted acinar cells, the levels of PAR6, one of the main components of the polarity complex that is directly activated by Cdc42 (Joberty et al.,2000) were significantly reduced (Fig. 2A), together with the levels of F-actin, which were not affected at the basolateral membrane (BLM) (Fig. 2A). The levels of ZO-1, one of the components of the TJ, were also reduced at the IC in Cre-expressing cells (Stevenson et al., 1986) (Fig. S2A) whereas, the levels and the organization of both E-cadherin, a marker of the adherens junctions (Fig. S2B), and non-muscle myosin IIA (Fig. S2C) were not perturbed. Next, we checked whether both cell polarity and structural integrity were maintained in the expanded IC. We found that the APM and the BLM did not mix, as shown by staining the acinar cells for the two well established salivary markers aquaporin 5 (AQP5) and the Na^+^-K^+^-Cl^-^ cotransporter 1 (NKCC1) (Fig. 2C) (Matsuzaki et al., 1999; O’Grady et al., 1987). Studies in other experimental systems have shown that downregulation of Cdc42 inhibits the formation of apical lumens and leads to the accumulation of large intracellular vacuoles (Kesavan et al., 2009; Martin-Belmonte et al., 2007; Sakamori et al., 2012). To confirm that the structures observed in salivary acinar cells were indeed derived from the ICs, we retro-injected low molecular weight fluorescent dextran into the main salivary duct in the anesthetized mice and imaged its delivery to the ICs by ISMic (Masedunskas et al., 2011a). We showed that in Cdc42-depleted cells, the enlarged ICs were still functionally connected to the main ductal system and, in addition, that paracellular integrity was not affected (Fig. 2D, S2D, and Movie S2).

**Figure 2.**
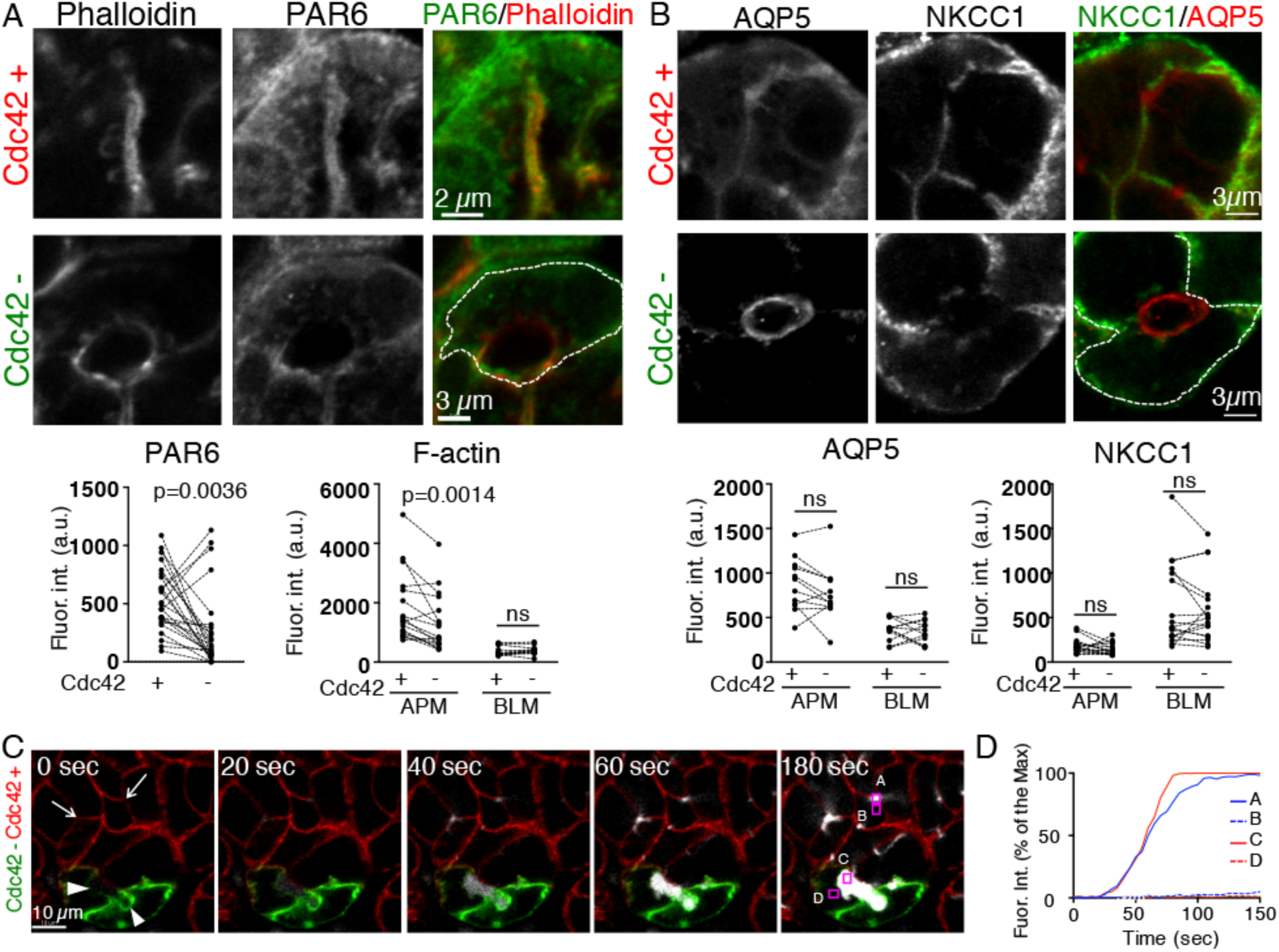
Cdc42-depletion induces loss of PAR6 and F-actin at the APM without altering polarity and junction permeability. **A-B**. Cdc42 ^fl/fl-mT/mG^ mice were transfected with Adeno-Cre and processed for immunofluorescence as described in legend to Figure 1. Samples were labeled with iFluor 405-Phalloidin (A, red) and an antibody against PAR6 (A, green) or with antibodies against AQP5 (B, red) and NKCC1 (B, green). Dotted-lines outline of Cre-expressing cells. Lower graphs shows the analysis of the levels of PAR6 (A), F-actin (A), AQP5 (B), and NKCC1 (B) in Cre-expressing (Cdc42-) and Cre-non expressing (Cdc42+) cells. (PAR6, N=25 ICs, 11 acinar cells from 3 animals; F-actin, N=17 APM and BLM, 12 acinar cells from 4 animals, AQP5, N=12 APM and BLM, 9 acinar cells from 3 animals; and NKCC1, N=17 APM and BLM, 17 acinar cells from 5 animals). ns=not significant, paired t-test. **C-D**. Three kDa Cascade Blue Dextran (White) was retro-diffused into the Wharton’s duct of anesthetized Cdc42 ^fl/fl-mT/mG^ mice after 3 weeks from Adeno-Cre transfection. **C.** The SMGs were exposed and imaged by ISMic. Time 0 represents the point at which was detected in IC. Arrow and arrowhead in time 0 shows normal IC and expanded-IC, respectively (C and Movie S2). **D.** Quantification of Dextran fluorescence intensity within (A, C) and outside (B, D) in Cre-negative cell (A, B) and Cre-positive cells (C, D). Graph shows a representative experiment.

### Membrane trafficking is altered in Cdc42-depleted cells

Based on the increase in the surface area of the ICs, we hypothesized that their expansion could be the results of an imbalance in membrane trafficking towards and from the APM. Consistent with this idea, we found that the cytoplasm of Cdc42-depleted cells contained GFP-labeled vesicular structures with apparent diameters varying from 0.5 to 2.5 μm. These vesicles were not detected in control cells (Fig. 3A, arrows, and Fig. 3B). Interestingly, we observed large GFP-containing vesicles in the subapical area which occasionally generated tubular structures (Fig. 3B, inset). Indirect immunofluorescence revealed that a subpopulation of vesicles smaller than 1.5 μm was endosomal in nature, as it was labeled by the early endosomal marker EEA1 (25 ± 6%, Av ± S.E.M N=3 animals, 44 cells, 401 vesicles) (Fig. 3C) and only occasionally with the small GTPase Rab11a (not shown), which has been previously shown to label the apical recycling endosomes and to control APM maintenance by regulating membrane trafficking from the Golgi apparatus and the early apical endosomes (Bai and Grant, 2015; Balklava et al., 2007; Winter et al., 2012). Depletion of Cdc42 did not alter the number, size, and cellular distribution of the EEA1-positive early endosomes (Fig. 3C) or the Rab11-positive compartments (not shown). The remaining GFP-labeled vesicles were not labeled by makers such as LAMP1 (Lysosomes), TGN46 (Trans-Golgi Network), GM130 (Golgi), VAMP4 (post-Golgi), or LC3 (autophagosomes) (not shown).

**Figure 3.**
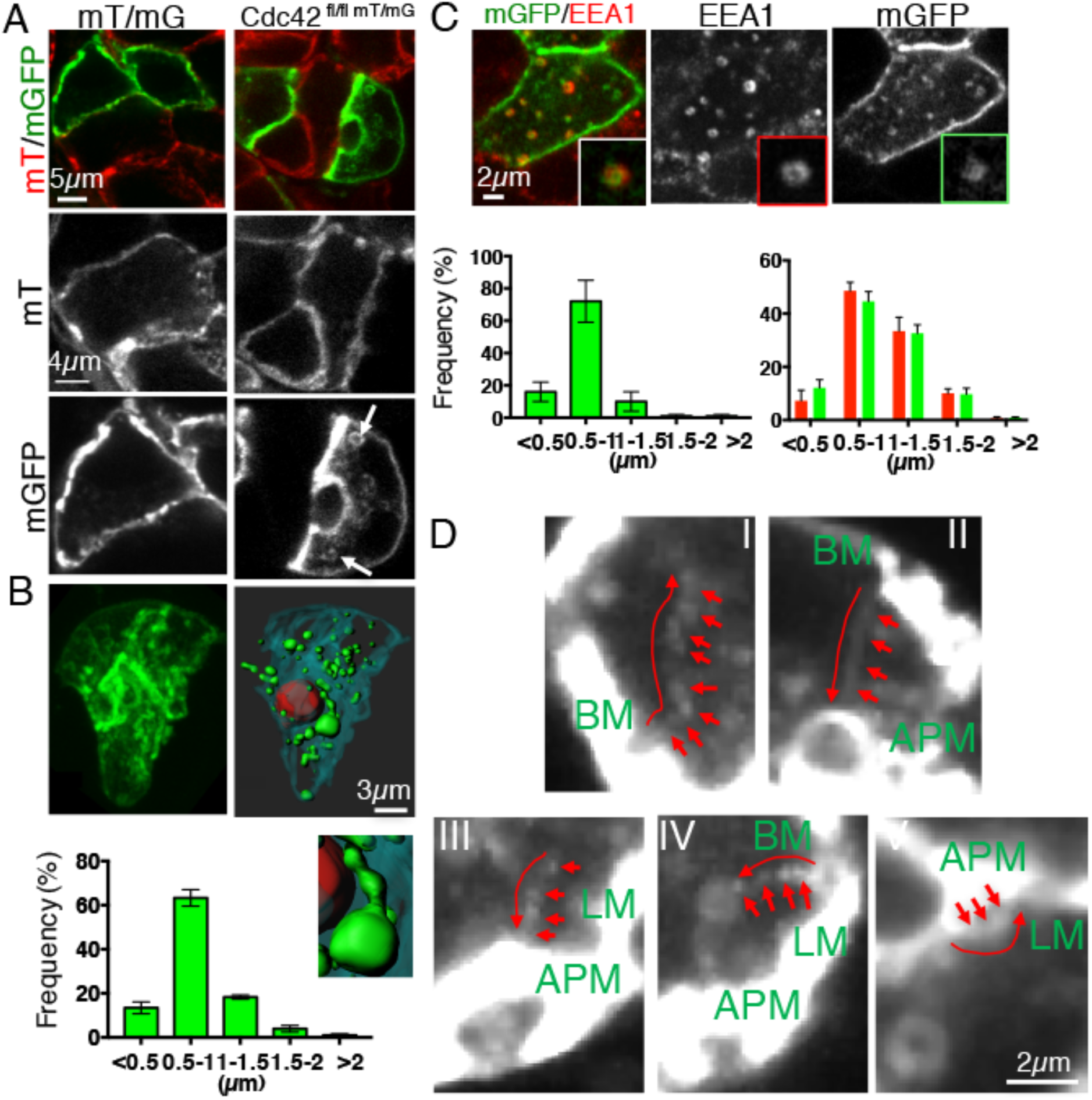
Membrane trafficking was altered in Cdc42-depleted cells. **A-D.** mT/mG (A, left panel) or Cdc42 ^fl/fl-mT/mG^ (A right panel, B-D) mice were transfected with Adeno-Cre, as described in Material and Methods. After 3 weeks, the glands were excised and processed for immunofluorescence (A-C) or imaged by ISMic after β-adrenergic stimulation (D). (A-C) Excised glands were left untreated (A-B) or labeled with antibodies against EEA1 (C). **A-B.** Confocal images of single sections (A, Arrows label mGFP positive vesicles) or Z-stacks (B, Cyan BLM, Red APM, green vesicles; inset shows a membranous tubule forming from a large vesicle. Graph in B shows the size distribution of the mGFP-positive vesicles in the cells (N=619 vesicles, in 4 animals, Data are Means ± S.E.M.). **C.** Co-localization between mGFP vesicles (green) and EEA1 (C, red). Quantification of the size distribution of mGFP vesicles which colocalize with EEA1 (left graph, N=351 vesicles in 3 animals) and the size distribution of EEA1 vesicles in Cre-positive (green bars) and Cre-Negative (red bars) (N=619 vesicles in 4 animals). Data are Means ± S.E.M.). **D.** ISMIc of Cre-positive cells. mGFP-labeled vesicles were tracked, as described in Material and Methods. Each panel shows an overlay of different time points (see Movie S3). Arrows point to the trajectory of selected vesicles which traffic from the basal membrane (BM) to the center of the cell (I), the BM to the APM (II), the center of the cell to the APM (III), the BM to a large vesicle (IV), and the APM to the lateral membrane (LM).

To reveal the direction of trafficking of the GFP-labeled membranes in Cdc42-depleted cells, we used ISMic. These vesicles were very dynamic (Movie S3) and they were transported towards several locations throughout the cells. We identified various patterns: 1) vesicles generated from the basolateral membranes and directed toward the center of cell (Fig. 3D, I) or the APM (Fig. 3D, II), 2) vesicles directed to the APM (3D, III);3) vesicles that fused or departed from large vesicles localized in the subapical areas (Fig. 3D, IV); and 4) vesicles generated from the APM and fusing with the lateral domains (Fig. 3D, V). These patterns were observed regardless of the cells were under basal conditions or stimulated to secrete proteins via β-adrenergic dependent regulated exocytosis. Due to the temporal and spatial limitation of light microscopy, we were not able to always visualize the initial steps in the internalization, and since the time-lapse images were acquired in a single focal plane, we could not track the vesicles for long distances, thus making it difficult to provide a quantitation of the frequency of these events. Nonetheless, these results strongly suggest that the expansion of the ICs could be linked to increased endocytic trafficking which delivers an excess of membranes to both the APM, and the lateral membranes.

Notably, we ruled out that the IC expansion was due to an imbalance between regulated exocytosis of the large secretory granules and the subsequent compensatory endocytosis (Masedunskas et al., 2011b). This conclusion was supported by two findings. First, Cdc42-depletion in acinar cells did not affect the β-adrenergic receptor-dependent fusion of the secretory granules with the IC, although it delayed their integration into the APM (Fig. S3A, S3B, and Movie S4); this finding is consistent with a reduction in the levels of F-actin recruited on the granules (Fig. S3C), which control this process *in vivo*, as we reported (Masedunskas et al., 2011a; Milberg et al., 2017). Second, the integration of the secretory granules did not result in a further expansion of the IC, indicating a rapid retrieval of the granular membranes via compensatory endocytosis (Sramkova et al., 2009) (Movie S4).

### Cdc42 depletion impairs the formation of IC postnatally

In other model systems, it has been shown that maintenance and formation of the epithelial lumen share common mechanisms (Joberty et al., 2000; Rojas et al., 2001; Suzuki and Ohno, 2006). Therefore, we investigated whether Cdc42 controls the development of the IC in salivary acinar cells and negatively regulates endocytosis under these conditions. To this end, we crossed the Cdc42 ^fl/fl-mT/mG^ mouse with a strain which expresses Cre under the control of the salivary gland-specific AQP5 promoter (ACID-Cre), which is activated in both acinar and intercalated duct cells at embryonic day 15 (Flodby et al., 2010) (Fig. S4A and S4B). Although the animals were viable, we measured a significant loss in body weight (Fig. S4C), and no significant differences were observed in the organization of salivary tissues (Fig. S4D). In the acinar cells of adult Cdc42-/-mice, the levels of Cdc42 were reduced by 50%, as shown by quantitative immunofluorescence (Fig. S4E).

First, we determined the kinetics of formation of the IC in acinar cells, a process that is completed after birth (Tucker, 2007). In control animals, central luminal structures were observed at post-natal day 1 (P1), whereas the IC were not developed yet (Fig. 4A). At P5 the IC began to sprout from the central lumen and to extend toward the basal membranes of the acini. At P15 they were fully developed and comparable to adult weaned mice (Fig. 4A and 4B upper panels, arrowheads, Movie S5). On the other hand, in Cdc42-depleted acini, the IC did not sprout from the central lumens. At P15 and in adult mice we observed one or two vacuolar-like structures within the same acinus (Fig. 4A and 4B lower panels, and Movie S5), similar to what previously reported (Kesavan et al., 2009; Martin-Belmonte et al., 2007; Sakamori et al., 2012). These represent, most likely, the final stage of the involution of the central lumens, as they were functionally connected to the ductal system (Fig. S4F, Movie S6). Electron microscopy revealed that in the adult Cdc42^fl/fl^-ACID-Cre mice the central luminal structures had reduced microvilli bulging into the lumen when compared to control mice (Fig. 4C), a phenotype that was reported in the liver of Cdc42-/-mice (van Hengel et al., 2008). Finally, we observed the progressive accumulation of mGFP-labeled vesicles that were not detected under control conditions (Fig. 4A lower panels, green arrowheads). Similar to what observed in the adult mice, these mGFP-labeled vesicles were labeled by EEA1 but not Rab11a (Fig. 4D and insets), thus indicating that the Cdc42 negatively regulate endocytosis during development, as well.

**Figure 4.**
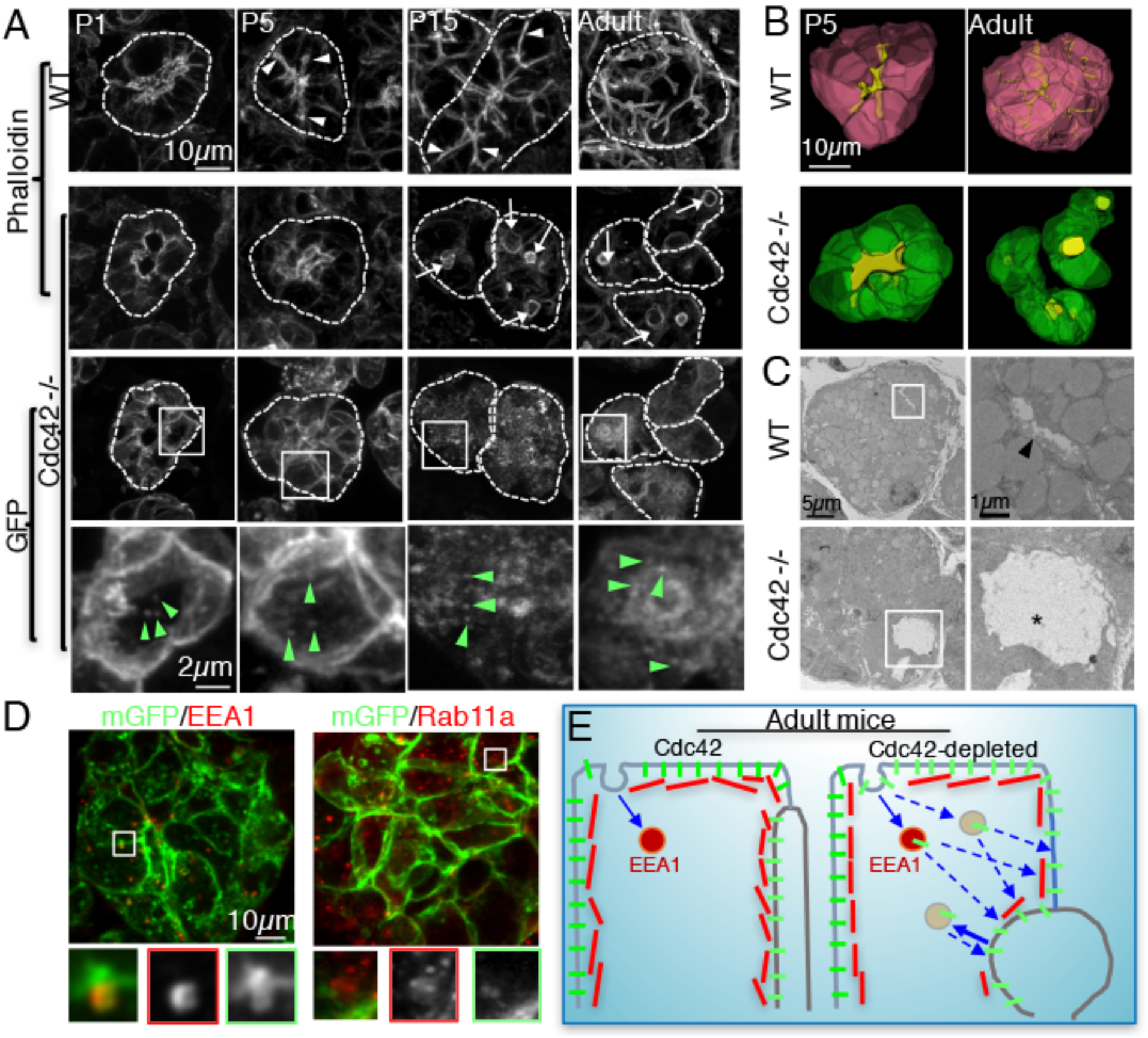
Cdc42 depletion impairs the formation of IC postnatally. **A-D**. The SMGs of either Cdc42 fl/fl-mT/mG (WT) or Cdc42 ^fl/fl-mT/mG^ /ACID-Cre mice (Cdc42 -/-) were excised at P1, P5, P15, and week 28 week (Adult). **A-B.** Samples were labeled with Phalloidin and Z-stacks were acquired by confocal microscopy, as described in Material and Methods. **A**. Maximal projections of the SMGs. Arrowheads and arrows highlight developed IC and involuted central canaliculi labeled with Phalloidin (upper panels). Lower panels show mGFP vesicles in the cytoplasm (insets, green arrowheads). Dotted-lines show the outline of acini. **B.** Volume rendering of acini. **C**. EM micrograph of the SMG acini of WT (upper panels) and Cdc42-/-mice (lower panels). Insets - The arrowhead and asterisk show an IC and lumen, respectively. **D.** SMGs of Cdc42 -/- mice were processed for immunofluorescence and labeled with antibodies directed against either EEA1 (left panels) or Rab11a (right panels). **E**. Proposed model

Here we show that *in vivo* Cdc42 plays a fundamental role in maintaining the apical lumen in adult mice and in controlling its post-natal development. Cdc42 has been already described to regulate these processes in other organs and in cell cultures (Kesavan et al., 2009; Martin-Belmonte et al., 2007; Sakamori et al., 2012; van Hengel et al., 2008) by controlling polarity complexes, actin cytoskeleton, and membrane trafficking. Not surprisingly we found that depletion of Cdc42 resulted in the loss of PAR6, as shown in mammalian epithelial cell culture and *Drosophila* epithelia (Harris and Tepass, 2010). However, whereas *in vitro* Cdc42 has been shown to enhance endocytic pathways via the regulation of the actin cytoskeleton (Georgiou et al., 2008; Leibfried et al., 2008; Sathe et al., 2018), in the acinar cells of salivary glands *in vivo* we found that it negatively regulates this process. Indeed, downregulation of Cdc42 induced a robust internalization of the membrane-targeted reporter mGFP, that is localized both at the basolateral and apical PM under control conditions. Resident basolateral, apical, or junctional proteins were not affected. In addition, we did not observe any gross alteration in the morphology of the early, late or recycling endosomal compartments. This raises the intriguing possibility that Cdc42, in addition to positively regulate membrane trafficking pathways from and to the APM, it may also prevent the internalization of selected proteins and lipids, which do not possess specific trafficking signals. Therefore, the enlarged canaliculi and the increase in the length of the apical-lateral border elicited by the downregulation of Cdc42 could be explained by an unbalance of trafficking events, which result in a net increase of the delivery of internalized membranes to both sites (Fig. 4E).

Whether this phenotype is related to a defect in the assembly of F-actin at the PM is still to be determined. Since cortical actin has been proposed to work as a functional barrier to prevent membrane fusion (Trifaro et al., 2008), reduced levels of F-actin at the IC could result in an increase in constitutive exocytic fusion events. This could account for the increase in the IC surface area. In addition, cortical actin has been shown to control various steps of endocytosis by regulating, for example, membrane tension at the PM (Boulant et al., 2011) or vesicle scission (Kaksonen et al., 2005; Romer et al., 2010). Although in yeast, F-actin positively regulates the internalization process, in mammals it is dependent on the cell type and the specific endocytic pathway (Goode et al., 2015; Hinze and Boucrot, 2018). Therefore, it is conceivable that in the salivary acinar cells *in vivo* inhibition of Cdc42-mediated F-actin assembly may simultaneously inhibit selected endocytic pathways and enhance others, as previously shown in *Drosophila* (Harris and Tepass, 2008). This applies to the APM of Cdc42-depleted acinar cells, where we detected reduced levels of F-actin. Vesicles internalized from the IC may be delivered to the adjacent lateral membrane, thus contributing to the increase in the size of the apical-lateral border. However, at the basal membrane Cdc42 may control endocytosis through a different mechanism, since the F-actin levels were not affected by Cdc42-depletion. Finally, when Cdc42 was ablated at late embryonic stages, the canaliculi did not form but the increase of endocytic activity was still observed, thus suggesting that Cdc42 negatively regulates endocytosis during development, as well.

In conclusion, this study reveals an additional role of Cdc42 in regulating membrane trafficking *in vivo* and underscores the value of ISMic in live rodents to investigate the dynamics of membrane remodeling under physiological conditions. Our findings will lead us to address several mechanistic questions on the relationship among signaling, actin cytoskeleton, and membrane trafficking during PM homeostasis *in vivo*. Specifically, we are poised to further define the nature of the Cdc42-dependent endocytic pathways implicated in this process and to elucidate the machinery operating downstream of Cdc42 both at the apical and the basolateral membrane.

## Author Contributions

A.S. and R.W. designed the experiments. A.S., L.M., S.E. performed the experiments. A.S., L.M., and D.C. analyzed the data. C.B. performed EM experiments. M.P.H. provided contributions with the developmental experiments. A.S. and R.W. wrote the manuscript. All authors read and approved the final manuscript.

## Acknowledgments

This research was supported by the Intramural Research Program of the NIH, National Cancer Institute, Center for Cancer Research and National Institute of Dental and Craniofacial Research. We would like to thank Dr. Julie Donaldson for critical reading of the manuscript; Drs. Zea Borok, Edward Crandall, and Peter Flodby for providing the ACID mice; Dr. James M. Anderson for providing the ZO-1 antibody; Cameron Keshavarz and Erin Stempinski for assistance during the EM preparation; and Dr. Yoshikatsu Sato and Dr. Yasuhiro Kamei for helping with the image processing.

## Materials and Methods

### Animals and procedures

All experiments were approved by the National Institute of Dental and Craniofacial Research (NIDCR, National Institute of Health, Bethesda, MD, USA) and National Cancer Institute (NCI, National Institute of Health, Bethesda, MD, USA) Animal Care and Use Committee. mT/mGFP mice were purchased from Jackson Laboratory (Bar Harbor, ME). Cdc42^fl/fl^, RhoA^fl/fl^, and Rac1^fl/fl^ mice were a generous gift of Dr. Yi Zheng (Cincinnati Children’s Hospital Medical Center, Ohio). ACID-Cre (AQP5-Cre) mice were a generous gift of Dr. Zea Borok (University of Southern California, California). These mice were crossed according to Fig. S1 and S4 and PCR was used to confirm their genotypes. All the mice (males and females) used in this study weighed 20–40 grams. Mice were anesthetized by an intraperitoneal injection of a mixture of ketamine (100 mg/kg) and xylazine (20 mg/kg).

### Adenovirus transfection into the mouse submandibular salivary glands

Adeno-Cre was prepared using the ViraPower Adenoviral Expression System as described in (Milberg et al., 2017) (Life Technologies, Carlsbad, CA). To inject the Adeno-Cre into the SMG, anesthetized mice were positioned in a custom-made device (Masedunskas et al., 2013) that allowed holding the mouth of the mice open under a stereomicroscope. A PE-8 cannula (Strategic Applications, Libertyville, IL) was connected to a 31G of sterile insulin syringe loaded with Adeno-Cre (~10^9^–10^10^ particles/gland in sterile saline) through a MicroFil Custom 35 gauge (World Precision Instruments, Sarasota, FL). The cannula was inserted in the main SMG duct (Wharton’s Duct) located below the tongue and stabilized with commercial Super-glue. 0.5mg/kg Atropine was injected sub-cutaneously 10 min before Adeno-Cre injection to prevent fluid secretion. The Adeno-Cre (total volume per gland, 20 μl) was injected using a PHD Ultra Nanomite syringe pump (Harvard Apparatus, Holliston, MA) at a flow rate of 5 μl/min. The cannula was removed after for 10 min from the injection. After recovery from anesthesia, mice were allowed to recover and placed back in their cages.

### Indirect immunofluorescence

Immunostaining for Adeno-Cre transfected gland and for ACID-Cre expressing gland was performed in non-frozen section and in cryo-sections, respectively. Primary and secondary antibodies used in this study are listed in Table 1. SMGs were fixed by cardiac perfusion using a solution consisting of 4% formaldehyde in 0.2 M HEPES buffered at a pH of 7.3, and post-fixed overnight. For non-frozen section, fixed glands were sliced using a vibratome (Leica, VT1000s, 150-200 μm thickness). For cryosections, glands were placed in optimum cutting temperature compound (Sakura Finetek USA Inc., Torrance, CA), snap-frozen in 2-methylbutane on liquid nitrogen, and cut using a cryostat (Leica, CM3050S, 10μm thickness). Immunostaining was performed as follows. Samples were incubated: 1) in 10% FBS and 0.02% Saponin in PBS (blocking solution) for 30-45 minutes at room temperature, 2) with primary antibodies in blocking solution at 4°C for either 2 days (non-frozen section) or overnight (cryo-sections), 3) with secondary antibodies in blocking solution at 4°C for either overnight (non-frozen section) or 30 min (cryosection), 4) if needed, with either Phalloidin or HOESCHT, for 30-60 minutes at room temperature. Finally, samples were mounted on a glass slide and covered with a #1.5 coverslips.

**Table 1.**
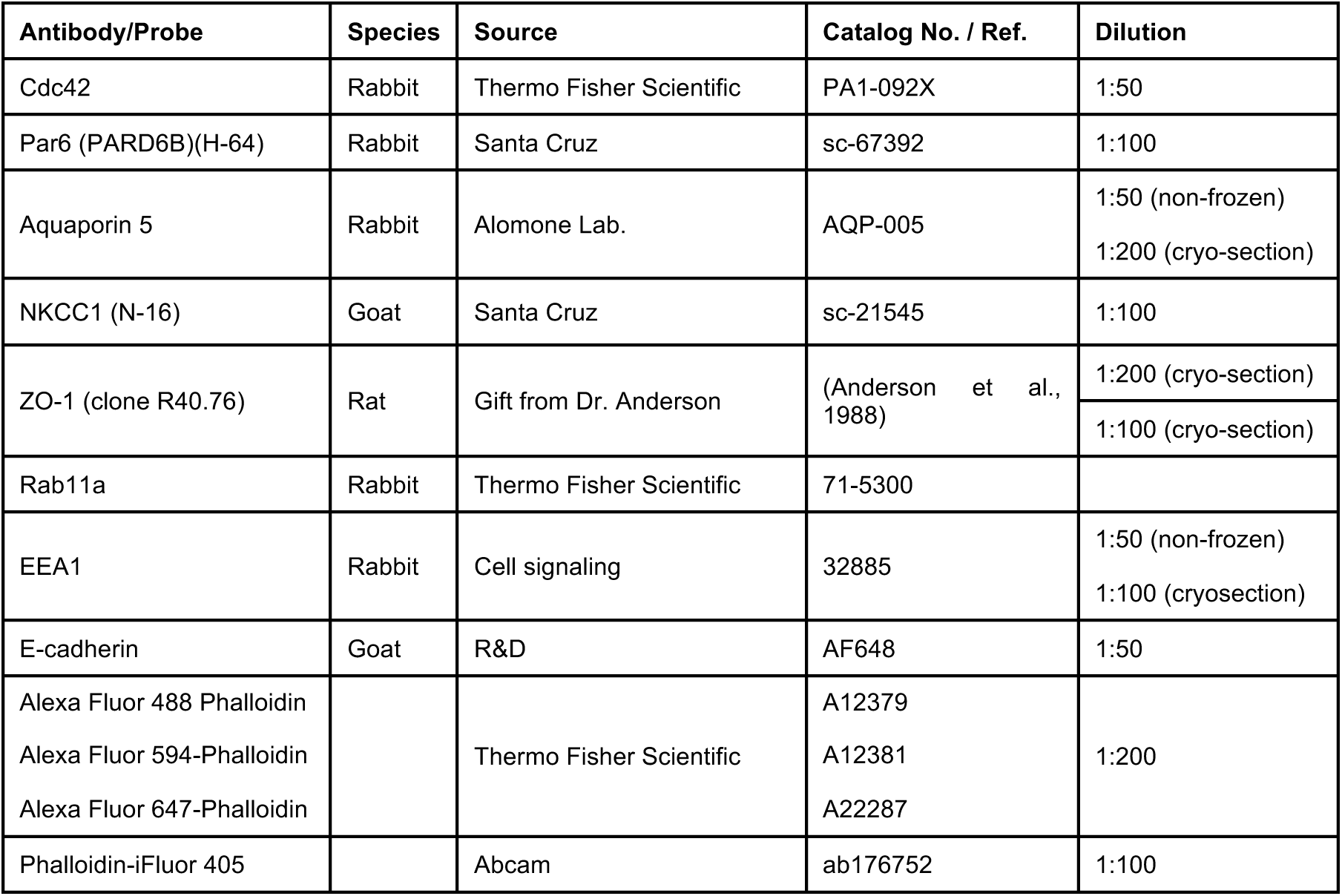
List of antibodies used in this study

### Intravital subcellular microscopy (ISMic)

In anesthetized mice, SMGs were exposed by a small longitudinal incision in the submandibular region. Connective tissue was separated from the glands without injuring the parenchyma, and the exposed glands were gently pulled out taking care of avoiding tissue damage. Mice were placed on the pre-heated stage of a confocal microscope (see next section) and covered with a heated pad (37–38 °C) to maintain the body temperature, as previously described (Masedunskas et al., 2013). The externalized SMGs were accommodated on a coverslip mounted on the stage above the objective and constantly moistened with a carbomer-940-based gel (Snowdrift Farm, Tucson, AZ) (Masedusnkas et al. 2008). The glands and the body of the animal were immobilized using custom-made holders, as previously shown (Masedunskas et al., 2011a).

To deliver 3 kDa Cascade Blue Dextran (Thermo Fisher Scientific, Waltham, MA), anesthetized mice were cannulated first, and the SMGs exposed, as described above. Mice were placed under the microscope and imaged while performing the injections with the pump (see above) at a flow rate of 300-500 nl/min.

### Microscope and imaging parameters

ISMic and indirect immunofluorescence were performed by a point-scanning IX81 inverted confocal microscope equipped with a Fluoview 1000 scanning head (Olympus America Inc.). All images were acquired using a Plan Apo 60x N.A. 1.42 oil immersion objective (Olympus America Inc.). Fluorophores were imaged using the appropriate lasers as required by their excitation spectra (laser excitation 405 nm, 488 nm, 561 nm or 633 nm). Fluorophores with slightly overlapping emission ranges were imaged using the “sequential” scanning mode to avoid bleed-through. The optimal focal plane for imaging the acinar cells was set at ~15 μm below the surface of the gland, as determined by visualization of the collagen capsule that surrounded the acinar cells. For ISMic of exocytosis and dextran delivery, the acquisition speed was set a 2 sec/frame and 10 sec/frame, respectively. And the pinhole was optimally set to 0.9 μm. Z-stacks were acquired with a step size of 0.50-1.00 μm During acquisition, XYZ drift was manually corrected.

### Tracking analysis

To enhance the visualization of the vesicles, the brightness of the original movie was adjusted so that the threshold was 50% of the maximum original brightness of the whole movie and the grayscale gamma was set to 0.6. To manually track the vesicles, a customized MATLAB script was executed to display the movie frame by frame with pseudo-color equal-brightness contour to emphasize the shape of the objects. The displayed frame allowed to click on the image to select the position of the object identified as a vesicle by eyes. After the selection, the script automatically recorded the selected position and switched the image to the next frame to select the position of the same vesicle in the next frame.

### Transmission electron microscopy

The gland tissue was excised and fixed for 90 min in 2% glutaraldehyde, 2% formaldehyde (Electron Microscopy Sciences, Hatfield PA), in 0.1 phosphate buffer (Ph 7.2), post-fixed in aqueous 1% osmium tetroxide, block stained with 1% uranyl acetate, dehydrated in graded ethanol solutions, and embedded in EMbed-812 (Electron Microscopy Sciences). Thin sections were stained with uranyl acetate, and lead citrate then examined on a JEM-1200EX (JEOL USA) transmission electron microscope (accelerating voltage 80 keV) equipped with an AMT 6 megapixel digital camera (Advanced Microscopy Techniques Corp).

### Image Analysis and Quantitation

For measurement of fluorescence intensity images were acquired by confocal microscopy using the same laser power and detectors settings for control and Cdc42-depleted cells. Images with the highest fluorescence intensity were selected from Z-stacks, and the contours of the apical membranes, basolateral membranes, and fused secretory granules were traced manually. Total fluorescence intensity was measured using Image J and normalized for the area of the structure. For Adeno-Cre transfected glands, fluorescence intensities of Cdc42-depleted were compared with those of neighboring control cells. For quantifications of Cdc42 levels, immunofluorescence levels in Cre-expressing cells were normalized to that of neighboring control cells. Statistical significance was calculated using the paired or unpaired Student’s t-test. Paired tests were used when IF was measured in cells within the same acinus. The volume rendering and measurement of surface area were performed with Imaris (Bitplane, Belfast, United Kingdom) using the isosurface tool. The structure of intercellular canaliculi and cell shape were traced according to fluorescence of phalloidin and of mT/mGFP, respectively. The time-lapse movies from ISMic were processed as described elsewhere (Milberg et al., 2017). Prior to quantification, motion artifacts created during ISMic was stabilized using the Stackreg plugin in Fiji (NIH, Bethesda, MD). In mice expressing the mT/mGFP reporter, the diameter of fused secretory granules was estimated from the circular profiles at the APM labeled by either mT or mGFP. Diameters were measured in time series from the frame the profiles of the granules were clearly visible. Images were analyzed and assembled in Fiji and Imaris. Data analysis was done in Prism (GraphPad, San Diego, CA) and Excel (Microsoft, Redmond, WA).

## Legends to Supplementary Movies

**Movie S1**–Volume rendering derived from a Z-stack phalloidin-stained acini. Cre-positive cells (green), Cre-negative cells (red), IC (yellow)

**Movie S2 –** Left: Time-lapse of 3 kDa Cascade Blue-dextran diffusion into the IC of adeno-Cre transfected Cdc42 ^fl/fl-mT/mG^ visualized ISMic (5 sec/frame). Right: Maximal projection of a Z-stack acquired 180 sec after the injection of the dextran. Cre-positive cells (green), Cre-negative cells (red), Dextran (White)

**Movie S3 –** ISMic of representative Cre-positive cells where mGFP-labeled vesicles are tracked as described in Material and Methods. Arrows highlight the trajectories of selected vesicles. Basal membrane (BM), apical Plasma Membrane APM (II), lateral membrane (LM). Frame rate 2s per frame.

**Movie S4 –** ISMic of regulated exocytosis of secretory granules in mT/mG (Cdc42+) and Cdc42 ^fl/fl-mT/mG^ (Cdc42 -/-) mice transfected with Adeno-Cre. Low magnification (Upper movies) and insets (lower movies) showing the integration of the secretory granules into the APM.

**Movie S5**–Volume rendering derived from a Z-stack of phalloidin-stained acini in SMGs excised at postnatal day 5 (P5, left) and week 28 (Adult, right) from mTmG mouse (Cdc42+) and Cdc42 ^fl/fl-mT/mG^/AQP5-Cre mice (Cdc42-). Cre-positive cells (green), Cre-negative cells (red), IC (yellow)

**Movie S6 – (related to Fig. S4)** –Retrograde delivery of 3 kDa Cascade Blue-Dextran into the IC of Cdc42 ^fl/fl-mT/mG^ /AQP5-Cre mice visualized by ISMic (2 sec/frame).

---

**Figure S1.**
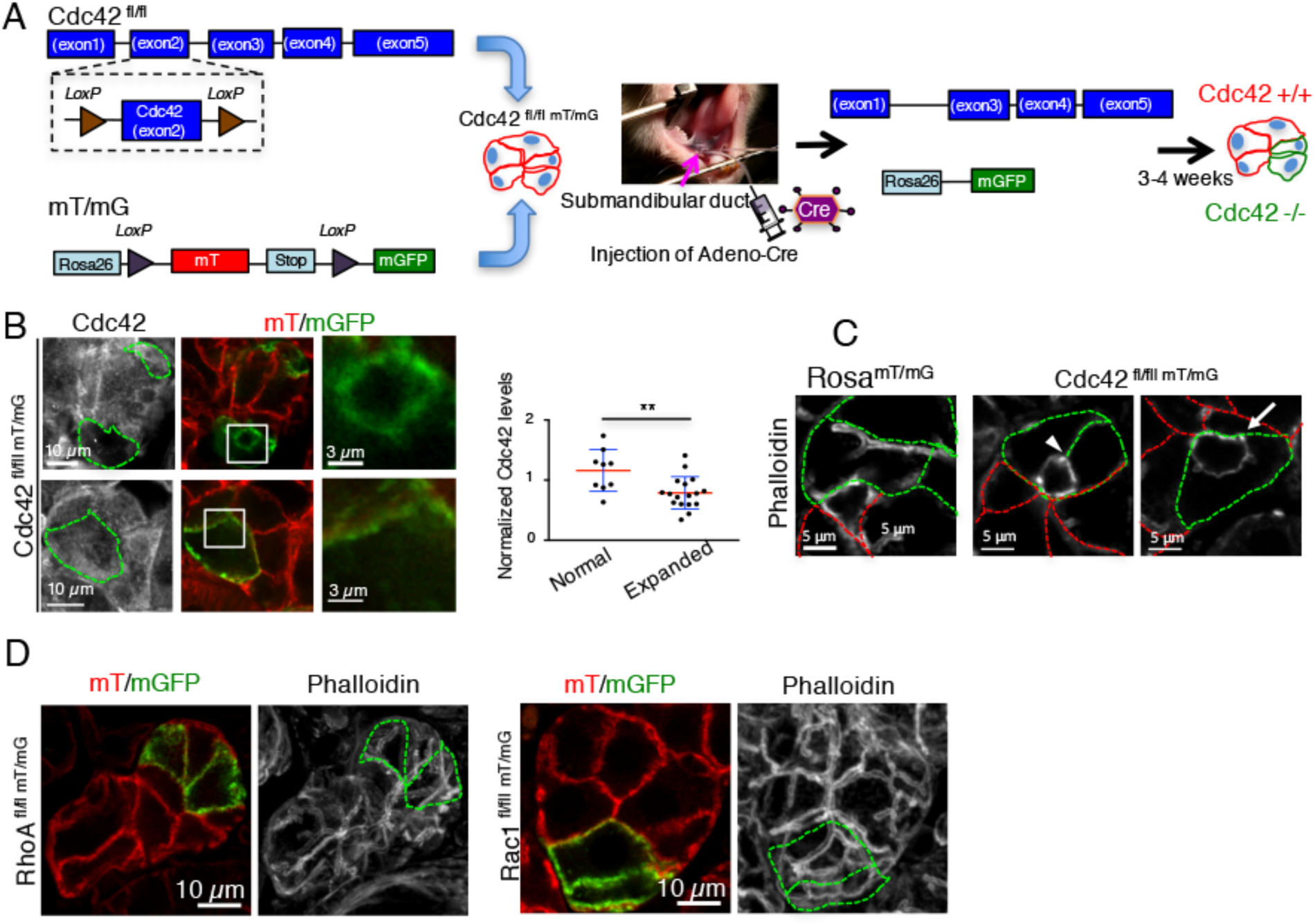
Characterization of the Cre/loxP-based mouse model for Cdc42 ablation. **A**. Scheme of the construction of the Cdc42 ^fl/fl-mT/mG^ mouse. Mice in which loxP sites flank the Exon2 of the Cdc42 gene were crossed with a strain expressing the Cre reporter Rosa^mT/mG^, in which a plasma membrane-targeted peptide tagged with the tandem-tomato protein (mT) is replaced by a similar peptide tagged with GFP (mGFP) upon Cre expression. Adeno-Cre is delivered to the SMG by retro-injection into the salivary ducts of anesthetized mice as described in Material and Methods. Experiments were performed 3-4 weeks after the adenovirus injection. **B,C**. The SMGs of Cdc42 ^fl/fl-mT/mG^ (B,C) or mT/mG (C) mice were excised after 3 weeks from Adeno-Cre expression, processed for immunofluorescence, and labeled with an antibody directed against Cdc42 (B) or with Alexa 647-Phalloidin (C). **B**. Correlation between the levels of Cdc42 and the phenotype of the IC. Left panels - Dotted-lines show the outlines of Cre-expressing cells and insets show high magnification of the expanded IC. The right graph shows the quantification of the levels of Cdc42 (determined by immunofluorescence, as described in Material and Methods) in Cre-expressing cells which have normal (1.16 ± 0.35, N=9) or expanded IC (0.79 ± 0.27, N=17 IC from 19 acinar cells in 3 animals). Data are Means ± S.D.; ** p<0.01, unpaired Student’s t-test. **C**. Green and red dotted lines outline Cre-positive and Cre-negative cells, respectively. Note that one cell lacking Cdc42 (arrows) is enough to alter the apical-lateral border (arrowhead). **D**. The SMGs of RhoA ^fl/fl-mT/mG^ and Rac1 ^fl/fl-mT/mG^ mice were excised 4 weeks after the retro-injection of Adeno-Cre into the salivary duct, processed for immunofluorescence, and stained with Alexa 647-Phalloidin. Left panels (mT/mGFP) represent single optical sections. Right panels (Phalloidin) are maximal projections of Z-stacks. Dotted-line show the outline of Cre-positive cells.

**Figure S2.**
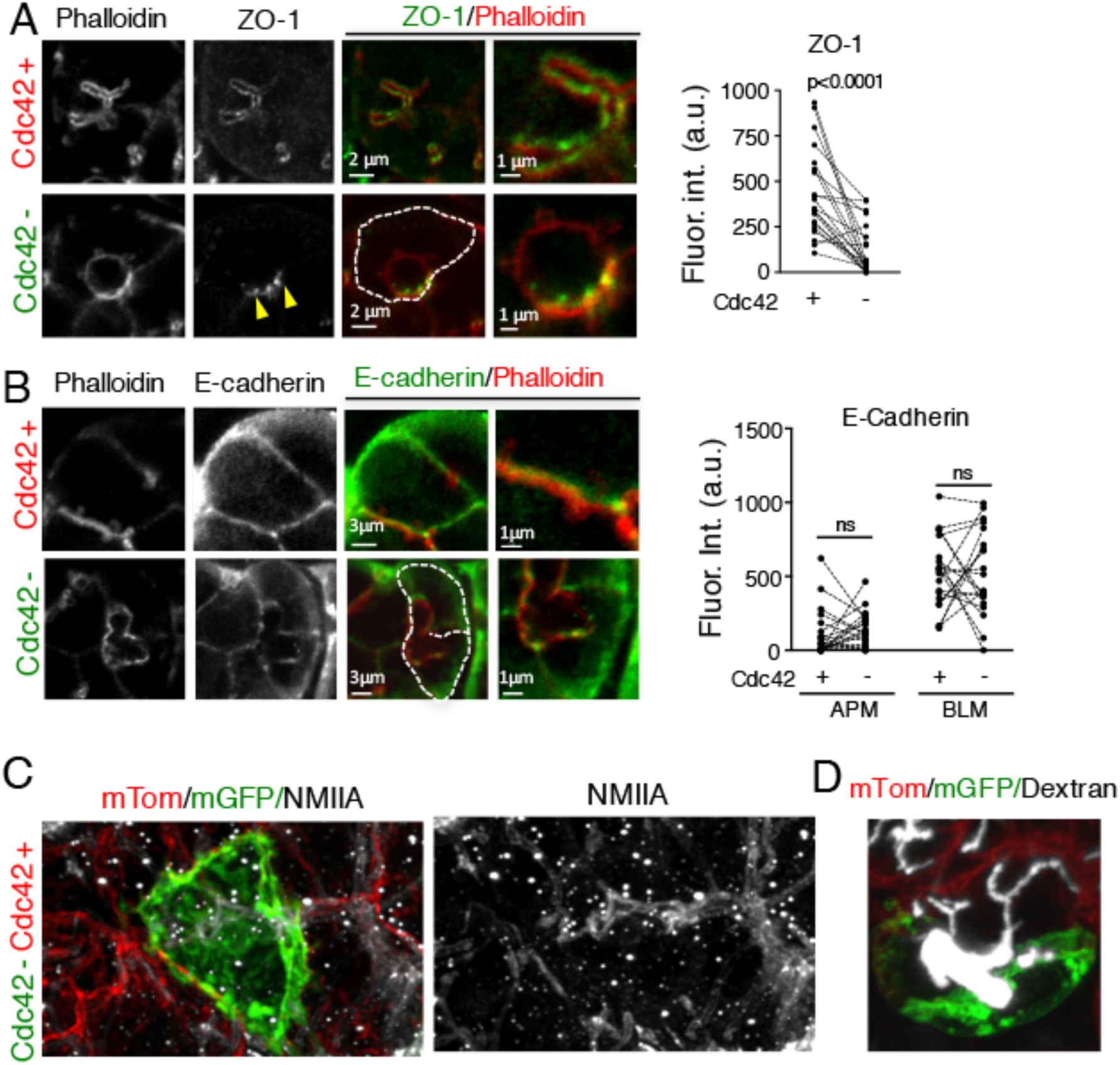
Characterization of the Cdc42-depletion in SMGs acinal cells. **A-D.** The SMGs of Cdc42 ^fl/fl-mT/mG^ mice were injected with Adeno-Cre, and either exposed and imaged by ISMic (D) or excised, processed for immunofluorescence, and labeled with primary antibodies directed against ZO-1 (A), E-Cadherin (B), NMIIA (C), and Phalloidin-iFluor 405 (A,B). **A,B.** Right panels show high magnifications of the APM. Dotted-lines show the outline of the Cre-expressing cells. Arrowheads in A show a cell-cell contact site. Right graphs show the analysis of the levels of ZO-1 (A) at the APM, and E-cadherin (B) at the ALM and BPM in Cre-expressing (Cdc42-) and Cre-non expressing (Cdc42+) cells. (ZO-1, N=21 ICs, 10 acinar cells from 4 animals; E-cadherin, N=20 APM and BLM, 9 acinar cells from 3 animals). ns=not significant, paired t-test. **C.** Maximal-projection of a Z-stack. **D**. Three kDa Cascade Blue Dextran was retro-injected in the SMGs. Maximal projection of a Z-stack acquired after 180s from the appearance of the dextran into the IC (from Figure 4, see Movie S2). Arrowhead shows the connection between the expanded and the normal IC.

**Figure S3.**
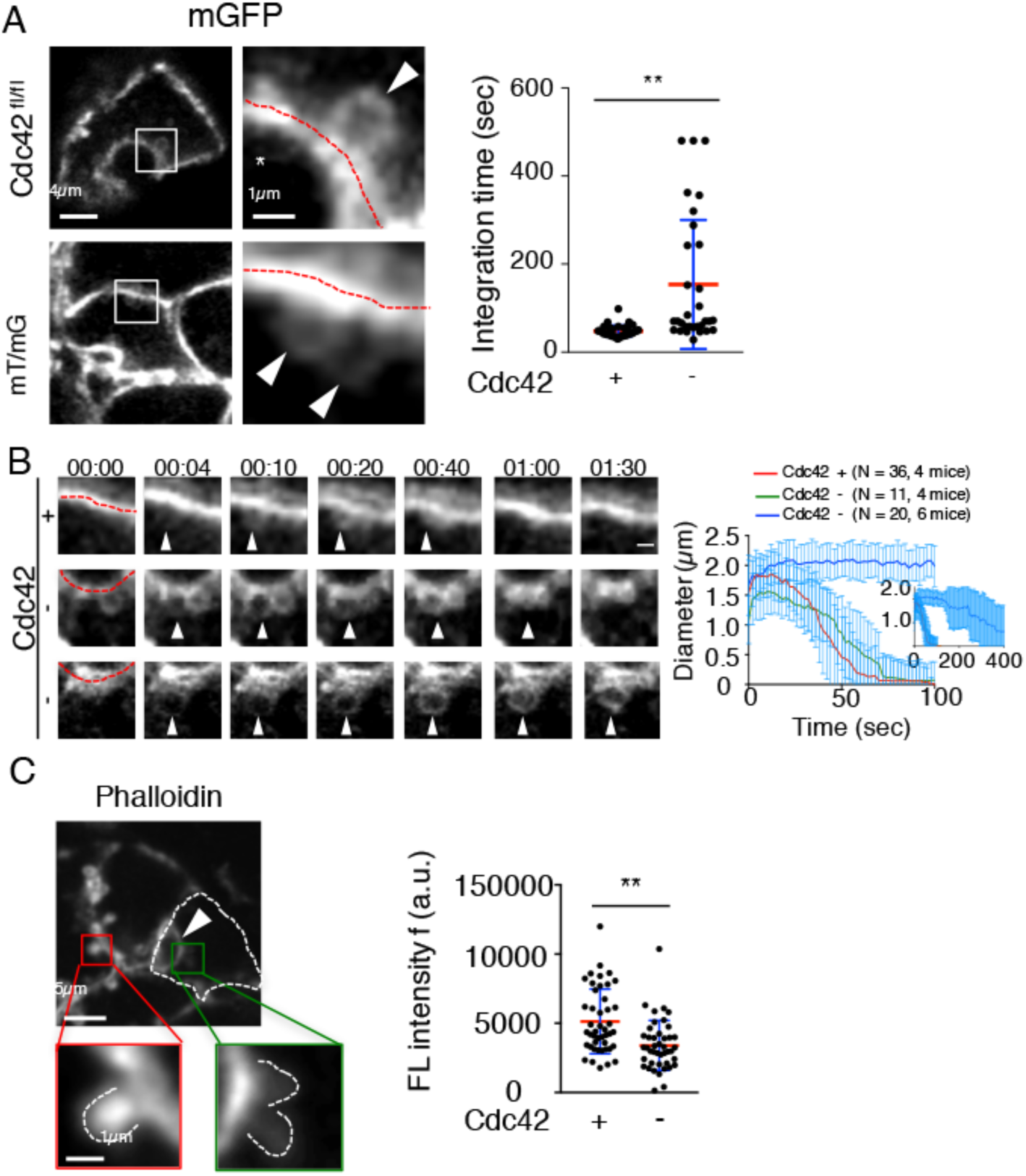
Cdc42 depletion delays the integration of the secretory granules during regulated exocytosis. **A-C**. Adeno-Cre was retroinjected in the Wharton’s duct of anesthetized Rosa^mT/mG^ (A-lower panels) and Cdc42 ^fl/fl-mT/mG^ (A-upper panels, B, C) mice. After 3 weeks, mice were injected sub-cutaneously with 0.02 mg/Kg isoproterenol (ISO), as previously described (Milberg et al., 2017). The SMGs were either exposed and imaged by ISMic (A-B) or excised and labeled with Alexa 647 Phalloidin (C). **A**. Snapshots of Cre-expressing cells (left panels) and insets showing secretory granules (SGs, left panels, arrowheads) fused at the APM (left panels, red dotted lines). The integration of the SGs is shown in Movie S4. Right graph shows the quantification of the SGs integration time in control cells (48 ± 13 sec) and Cdc42-depleted cells (153 ± 146 sec), measured as described in Material and Methods. Data are Means ± S.D. **p<0.01, unpaired t-test (N=36 SGs from Cre-expressing cells in 4 mT/mG mice; N=31 SGs from Cre-expressing cells in 6 Cdc42 ^fl/fl-mT/mG^ mice). **B**. Time-lapse series of the integration of the SGs into the APM in Cre-negative (upper panel, +) and Cre-positive (middle and lower panel, -) cells. Arrowheads and red dotted-lines show SGs and APM respectively. Time 0 represents the point at which the limiting membranes of the SGs were detected. Right graph shows time-dependent changes of a diameter of SGs after fusion with the APM, and the inset graph shows a longer time scale of the same graph. Data are Means ± S.D., N=number of SGs from 4-6 animals. **C.** Depletion of Cdc42 causes the reduction of F-actin level around fused SGs. Phalloidin staining of Iso-stimulated Adeno-Cre transfected Cdc42 ^fl/fl-mT/mG^ SMG acinar cells. Dotted lines and arrowheads show the outline of Cdc42-depleted cells and expanded IC, respectively. The graph on the right shows the quantitative analysis of the F-actin levels on the fused secretory granules in control cells (N=45 SGs from 5 mice) and in Cdc42-depleted cells (N=43 SGs from 5 mice). Data are Means ± S.D. ** p<0.01, unpaired Student’s t-test.

**Figure S4.**
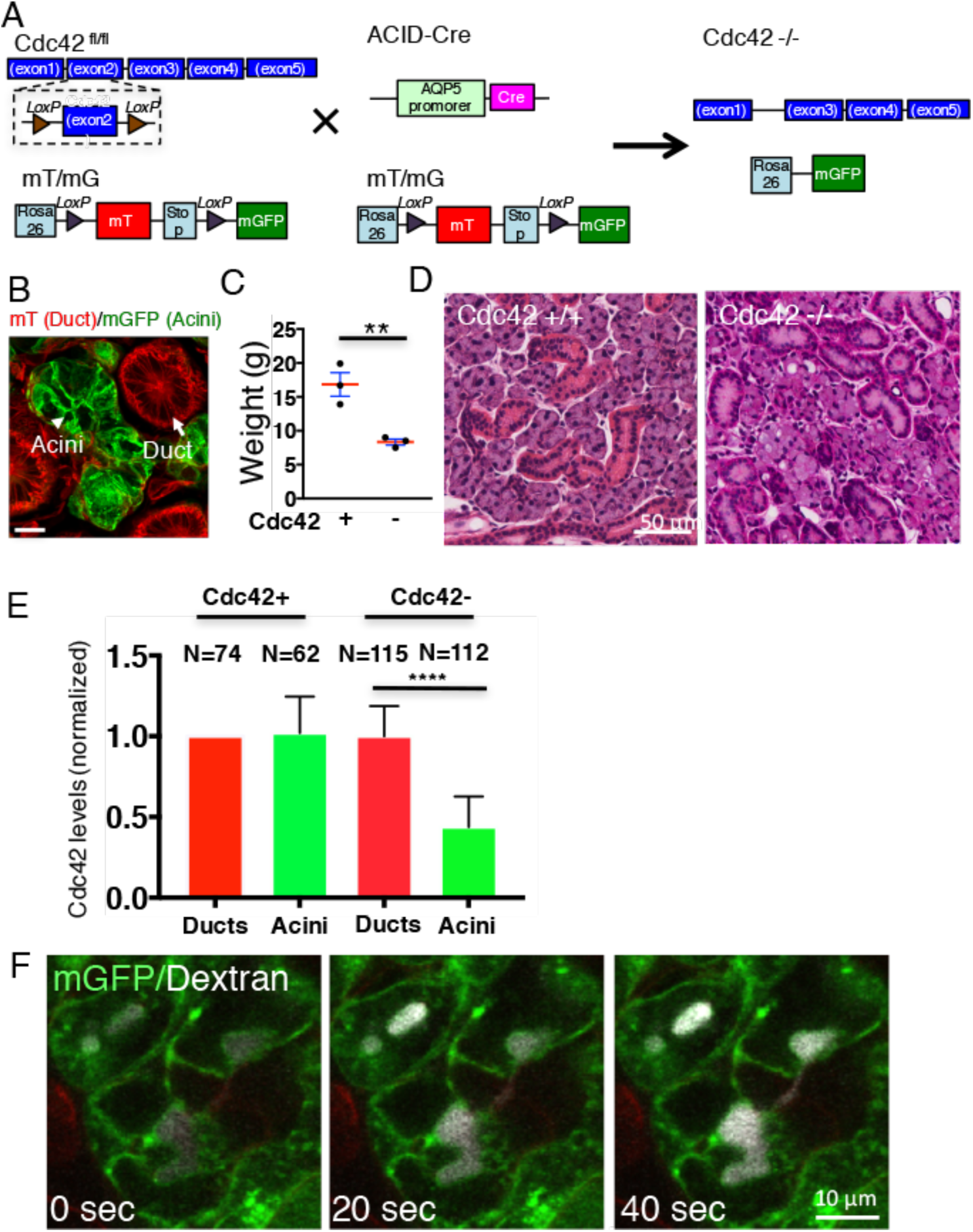
Characterization of late embryonic depletion of Cdc42. **A**. Cdc42 ^fl/fl-mT/mG^ mice were crossed with mice expressing Cre under the control of the AQP5 promoter (Cdc42 ^fl/fl^ ACID-Cre) in salivary acinar cells and intercalated ducts. Both strains were crossed with Rosa^mT/mG^ Cre reporter mouse. B. The SMGs were excised from mice at postnatal day 5 and processed for immunofluorescence. Arrow and arrowhead point to duct and acini, respectively. **C**. Weight of adult (28 weeks) mT/mG-ACID-Cre (Cdc42+) and Cdc42 ^fl/fl^ ACID-Cre (Cdc42-) mice. Data are Means ± S.E.M. (N=3 animals) **D**. SMGs were excised from adult mTmG/ACID-Cre (upper panel) and Cdc42 ^fl/fl^ /ACID-Cre mice (lower panel) and stained with Hematoxylin and Eosin. **E**-Levels of Cdc42 were measured in acinar (red bars) and ductal (green bars) cells in both mT/mG (Cdc42+) and Cdc42 ^fl/fl^ ACID-Cre (Cdc42-) mice at P15, as described in Material and Methods. Data which are normalized for the ductal cells in Cdc42+ mice are expressed as Means ± S.D. N represent the number of cells in one representative animal. **F**. ISMic of the retrograde delivery of 3 kDa Cascade Blue Dextran (White) into the central lumens of Cdc42 ^fl/fl^ -ACID-Cre mouse SMG. Time 0 represents the point at which the Dextran fluorescence was detected in the lumens (Movie S6).

